# Comparative study on the virulence of mycobacteriophages

**DOI:** 10.1101/2024.10.23.619922

**Authors:** Ilaria Rubino, Carlos A. Guerrero-Bustamante, Melissa Harrison, Sheila Co, Isobel Tetreau, Mani Ordoubadi, Sasha E. Larsen, Rhea N. Coler, Reinhard Vehring, Graham F. Hatfull, Dominic Sauvageau

## Abstract

The global tuberculosis (TB) epidemic affected 10 million people and caused 1.3 million deaths in 2022 alone. Multidrug-resistant TB is successfully treated in less than 60% of cases by long, expensive and aggressive treatments. Mycobacteriophages, viruses that can infect bacteria such as *Mycobacterium tuberculosis*—the species responsible for TB, have the potential to redefine TB prevention and treatments. However, the development of phage-based products necessitates the assessment of numerous parameters, including virulence and adsorption, to ensure their performance and quality. In this work, we characterized the virulence of three different mycobacteriophages (Fionnbharth, Muddy and D29), alone and as cocktails, against a TB model host (*Mycobacterium smegmatis*) under planktonic and early-stage biofilm growth conditions. Phage D29 and cocktails containing D29 had the highest virulence under all conditions. Interestingly, phages Fionnbharth and Muddy and their combination showed higher virulence against early-stage biofilm than against the planktonic phenotype. Adsorption assays indicated that all three phages had lower adsorption efficiencies on the early-stage biofilm phenotype than on the planktonic one, suggesting a reduced availability of receptors in the former. Given that, despite these lower adsorption efficiencies, the virulence of the phages and phage cocktails was either unchanged or higher against the early-stage biofilm, this phenotype must display properties that are favorable to other steps of the infection process. These results inform us on the dynamics of mycobacteriophage infections, alone and in cocktail formulations, under different host growth conditions, and serve as a basis for the development of phage products targeting mycobacteria biofilms.

## 1. Introduction

Tackling the tuberculosis (TB) epidemic, caused by *Mycobacterium tuberculosis*^1^, is a primary global health challenge. TB is among the top 10 causes of death in the world and has the highest mortality rate among infectious diseases; it infected 10 million people and caused 1.3 million deaths in 2022 alone^2^, and rising drug resistance limits current treatments^3^. For example, multidrug-resistant TB (MDR-TB)—caused by strains not responding to the main antibiotics used in treatment—is successfully treated by second-line drugs in only 63% of cases^2^. Compared to first-line drugs, second-line drugs have severe side effects^4^, demand even longer treatments (up to 2 years vs. 6 months), and are expensive^2^. Long therapies and severe side effects lead to poor regimen adherence. Additionally, extensively drug-resistant TB (XDR-TB), for which even second-line treatments fail, occurs in 15% of drug-resistant cases^2^. While efforts are ongoing to develop prophylactic, post-exposure, and therapeutic (against reoccurrence) vaccines, the only TB vaccine currently available is the bacille Calmette–Guérin (BCG). Inactivated/live whole cell, protein subunit, and viral vectored candidates are in clinical trials^5^, with M72/AS01E regarded as most promising (54% efficacy)^6,7^. Despite the merit of BCG in protecting children^8^ and other recent developments, no vaccine protecting adults has yet been approved.

Bacteriophages (phages)—viruses that infect bacteria—have great potential for several applications in health and other fields^9–20^. Phage-based technologies, in general, exploit the phage’s ability to recognize and lyse specific bacteria. Among several advantages of phage therapy^21^, the reduced rate in developing resistance, as compared to antibiotics, is particularly beneficial for TB. As such, mycobacteriophages, phages that target mycobacteria, have the potential to redefine prophylaxis, treatments and diagnostics for TB as well as for other mycobacterial infections^22–27^. Phages Fionnbharth, Muddy and D29 are examples of mycobacteriophages with high therapeutical potential; all three can infect *M. tuberculosis*, its commonly used low-containment model bacterium *Mycobacterium smegmatis*, and other hard-to-treat mycobacterial hosts^22,26,27^. For instance, Muddy (alone or in phage cocktails) has shown promising results in clinical cases of *Mycobacterium abscessus*^28–30^, *Mycobacterium chelonae*^30,31^ and *Mycobacterium avium*^30^ infections. A patient with BCG mycobacterial infection partially responded to a phage cocktail including Fionnbharth, Muddy and D29^30^; and a patient with *M. abscessus* showed favorable clinical and microbiological responses to a cocktail containing D29^30^. A 5-phage cocktail (including Fionnbharth, Muddy and D29) has been proposed for the treatment of TB^26^. Additionally, nebulized D29 has been shown to provide some prophylactic prevention of *M. tuberculosis* infection in mice^32^.

To reliably assess and improve the performance and quality of mycobacteriophage-based products, it is important to evaluate, among other parameters, the efficacy of a phage or phage cocktail at infecting a host population under therapeutically relevant conditions. Although the interactions of phages with *Mycobacterium* are often evaluated using shaken planktonic cultures, static and biofilm cultures are also important phenotypes for therapeutic applications, as they more closely resemble the environment encountered by mycobacteriophages during mycobacterial infection (e.g., hypoxic, low-nutrient environment with reduced to arrested growth rate)^33^ and they are thought to favor antibiotic resistance by contributing to the process of caseous necrosis in these infections^34^. To this aim, we characterized the virulence of three clinically relevant mycobacteriophages (Fionnbharth, Muddy and D29) against the TB model host *M. smegmatis* under planktonic and early-stage biofilm conditions using the virulence index^35^ adapted for mycobacterial cultures. This was done for the phages alone and in combination. Additionally, given the importance of adsorption dynamics and efficiency on the infection process, the adsorption behavior of these three phages was characterized under similar conditions. These results serve as a base for the development of formulations and conditions for efficient phage-based treatments of mycobacterial infections.

## 2. Materials and Methods

### 2.1 Host and mycobacteriophages

The host was *Mycobacterium smegmatis* mc^2^155, grown in 7H9 medium (Middlebrook 7H9 broth supplemented with 10% (v/v) Middlebrook ADC, 0.2% (v/v) glycerol and 1 mM CaCl_2_) at 37 °C and 250 rpm (Ecotron shaker incubator; Infors HT, Anjou, Quebec, Canada) and maintained on 7H10 agar plates (Middlebrook 7H10 agar with 0.5% (v/v) glycerol). The colony forming units/mL (CFU/mL) were determined by incubating 10 μL of 10× dilutions onto 7H10 agar plates at 37 °C for 3 days (Fisherbrand Isotemp 500 series oven; Fisher Scientific, Ottawa, Ontario, Canada).

Protocols for the amplification of phages in liquid cultures were adapted for the amplification of mycobacteriophages FionnbharthΔ*45*Δ*47* – a genetically engineered phage derived from a temperate phage with integrase and repressor genes deleted to render it strictly lytic^36^ –, Muddy HRM^N0157^–2 – a host range mutant able to efficiently infect *M. tuberculosis*^26^ –, and D29 – a lytic derivative of a temperate phage^27^, as follows. A culture of *M. smegmatis* was grown to early exponential phase (optical density at 600 nm (OD_600_) ∼ 0.2–0.4; Ultrospec 50 UV/Vis spectrophotometer; Biochrom, Holliston, MA, USA). High-titer phage stocks were diluted in phage buffer (10 mM Tris pH 7.5, 10 mM MgSO_4_, 68.5 mM NaCl, 1 mM CaCl_2_) and used to infect *M. smegmatis* cultures at multiplicities of infection (MOIs) of 1 for Fionnbharth and Muddy and of 0.01 for D29. Following incubation (37 °C and 250 rpm) and OD_600_ dropping to ≤ 0.4 (Fionnbharth and Muddy: 24–26 h; D29: 7 h), the phages were purified by centrifugation (17,000 g and 20 °C for 5 min; Eppendorf 5424 Microcentrifuge; Eppendorf, Mississauga, ON, Canada) and filtration (0.2-µm syringe filters or vacuum filtration system (Corning, Corning, NY, USA), based on the volume processed). The range of titers obtained was 5×10^10^–1×10^11^ plaque forming units/mL (PFU/mL). The stocks were stored at 4 °C; for Fionnbharth and Muddy stocks, the 7H9 medium was replaced with phage buffer by using Amicon Ultra Centrifugal Filters with 100-kDA molecular weight cut-off (Millipore Sigma, Oakville, Ontario, Canada) and centrifuging (1,750 g at room temperature for up to 30 min; IEC Clinical Centrifuge; Needham, MA, USA).

The phage stocks were titered by spot testing, following well-established protocols^37^. Briefly, the bacteria lawn was obtained by mixing 500 µL of a 2-day old *M. smegmatis* culture with 4.5 mL of 7H9 top agar (7H9 medium with 0.7% (m/v) agar) and distributing homogeneously on 7H10 agar plates. After solidification, 10 µL of 10× dilutions of phage suspension, prepared in phage buffer, were spotted and the plates were incubated overnight at 37 °C. Titers were determined in triplicates or more.

Agar, Middlebrook 7H9 broth, Middlebrook ADC, and Middlebrook 7H10 agar were supplied by BD (Mississauga, Ontario, Canada); all other chemicals were supplied by Fisher Scientific.

### 2.2 Mycobacteriophage virulence

A schematic of the experimental procedures is shown in Figure 1A. The virulence index^35^ method was adapted for mycobacteriophages. To obtain the reduction curves (OD_600_ as a function of time; Supplementary Figure 1 and Supplementary Figure 2), *M. smegmatis* cultures were grown to 10^7^ CFU/mL, and infected with Fionnbharth, Muddy or D29 at MOIs ranging from 10^-7^ to 100, in pre-warmed 96-well plates (final volume: 220 μL). For phage combinations, two- and three-phage cocktails (equal proportion per phage) were used to obtain combined MOIs of 10^-7^–100. Incubation was carried out at 30 °C in a plate reader (BioTek Cytation 5; Agilent, Santa Clara, CA, USA) and the data were recorded using Biotek Gen5 software (Agilent); OD_600_ measurements were obtained every 1 h until the control cultures (no phage) reached stationary phase. The virulence assays were conducted under two growth conditions. In the first one, linear shaking at 410 rpm was applied (planktonic growth). In the second one, shaking was only applied for 10 s before each OD_600_ measurement; morphological appearance of the control (no phage) samples corresponded to biofilm pellicle formation observed at the liquid-air interface in early-stage biofilms for *M. smegmatis*^38^ and *M. tuberculosis*^39^.

**Figure 1.**
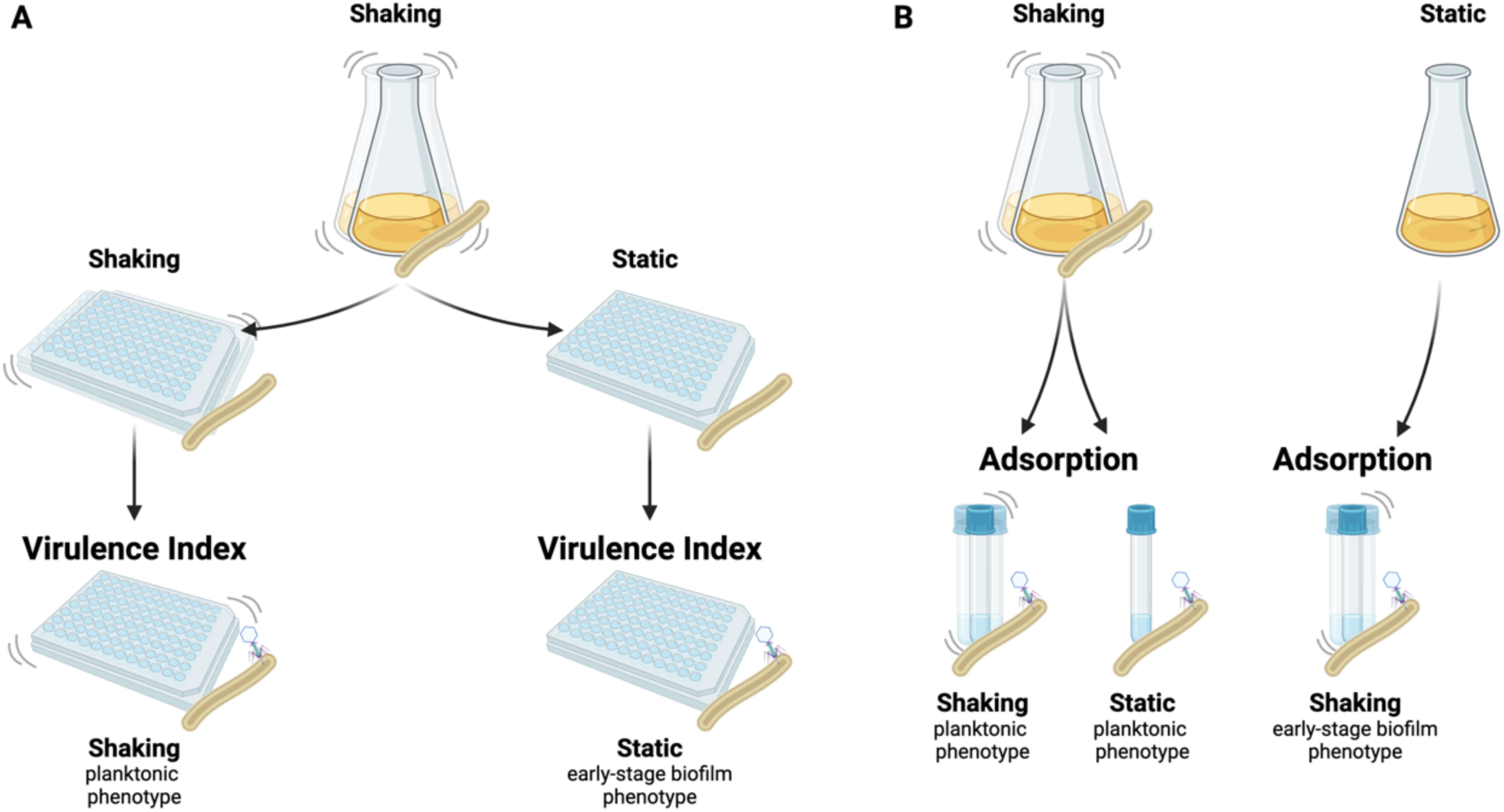
Schematic representation of the virulence and adsorption assays under different growth and assay conditions. Virulence (A): A planktonic *M. smegmatis* culture was transferred to 96-well plates to establish the planktonic (shaking growth condition) and early-stage biofilm (static growth condition) host cultures. Phages were added to perform the adapted virulence index protocol for each phenotype. Adsorption (B): Planktonic cultures (shaking growth condition) were used as the host for the adsorption assay performed under shaking and static conditions. Early-stage biofilm cultures (static growth) were used as host for the adsorption assay under shaking conditions.

The data was analyzed as reported previously^35,40^. Briefly, to obtain the local virulence, the areas under the reduction curves were calculated for each well, between the time of infection (time 0) and the time at which the no-phage control entered the stationary phase (determined as the time at which the slope of the OD_600_ over time was ≤ 0.03 h^-1^). The local virulence for a given MOI (ϖ_i_), describing the phage infection dynamics, was calculated as *v_i_* = 1 − *A_i_*⁄*A_0_*, where A_0_ and A_i_ are the areas under the OD_600_ curves for the no-phage control and for the infection at the MOI of interest, respectively. The virulence curve was obtained by plotting the ϖ_i_ values as a function of the log (MOI); the area under the virulence curve was normalized to obtain the virulence index (V_P_), which provides a score between 0 and 1 to quantify the virulence of a given phage or phage cocktail. At high MOI (100), under early-stage biofilm growth conditions, the three-phage cocktail started to show bacterial resistance before the end of the assay; in the analysis of the reduction curves at this specific condition, the OD_600_ value at 30 h incubation (before resistance was detected) was used thereafter.

### 2.3 Mycobacteriophage adsorption experiments

A schematic of the experimental procedures is shown in Figure 1B. Tests were conducted by adapting established protocols for phage adsorption^41^ for mycobacteria. For adsorption to planktonic *M. smegmatis*, a culture was grown to OD_600_ of 1 at 37 °C and 250 rpm. High-titer phage stocks were diluted to 1×10^9^ PFU/mL in 7H9 medium; 10 mL of host were infected at an MOI of 0.1. Immediately following infection, the samples were incubated at 37 °C, either at 0 rpm (static adsorption condition) or 250 rpm (shaking adsorption condition). Samples of the free phages were taken by transferring 10 μL into 1 mL of pre-chilled 7H9 medium to interrupt adsorption, followed by centrifugation (16,000 g, 4 °C, 5 min). The supernatants from each time point were titered to determine the free phage titer (P). The initial phage titer (P_0_) was determined similarly, but with 7H9 medium instead of host culture and without any incubation step. For adsorption to *M. smegmatis* early-stage biofilms, no shaking was applied during growth; the adsorption tests were carried out under shaking at 250 rpm, at 37 °C. The cell density of cultures used for the adsorption was 1.6 g/L and 1.7 g/L for the planktonic and early biofilm conditions, respectively.

The adsorption efficiency (ε) and rate constant × initial host concentration (*k·C_h0_*) were calculated by fitting (minimization of the sum of squared errors) the experimental data to the following model suggested by Storms *et al.*^42^:

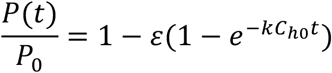

### 2.4 Statistical analysis

The statistical analysis was performed by using one-way analysis of variance (ANOVA), General Linear Model, and two-sample t-test using Minitab (Minitab, State College, PA). The values compared were considered statistically significant for P values less than 0.05.

## 3. Results and Discussion

### 3.1 Comparison of virulence of mycobacteriophages and cocktails

The virulence indexes of individual phages, 2-phage cocktails and the 3-phage cocktail infecting planktonic cultures are reported in Figure 2A (the corresponding virulence curves are found in Figure 3, and the respective reduction curves are found in Supplementary Figure 1). In planktonic cultures, phage D29 showed the highest virulence against *M. smegmatis*, when compared with phages Fionnbharth and Muddy (one-way ANOVA, P < 0.001, respectively; Figure 3H). Phage D29 exhibited high virulence even at low MOIs, resulting in a high virulence index value (V_D29-planktonic_ = 0.79; Figure 2A). Phages Fionnbharth and Muddy caused measurable virulence starting from MOI 10^-4^ and 10^-1^, respectively (Figure 3H). This resulted in low virulence index values (V_Fionn-planktonic_ = 0.20 and V_Muddy-planktonic_ = 0.14; Figure 2A). The combination of Fionnbharth and Muddy showed a purely additive effect on virulence, with no signs of synergistic or antagonistic effects (Figure 3H). All two-phage and three-phage cocktails containing D29 showed similar infection kinetics (General Linear Model, P > 0.05; Figure 3H) and virulence index values (one-way ANOVA, P > 0.05; Figure 2A) to D29 used alone. This was observed even at MOIs at which Fionnbharth and Muddy didn’t cause significant bacteria reduction. This is likely due to the high virulence of D29 and the absence of antagonistic effects from Fionnbharth and Muddy, which could allow it to overtake the infection process rapidly, achieving similar levels of virulence as when used alone.

**Figure 2.**
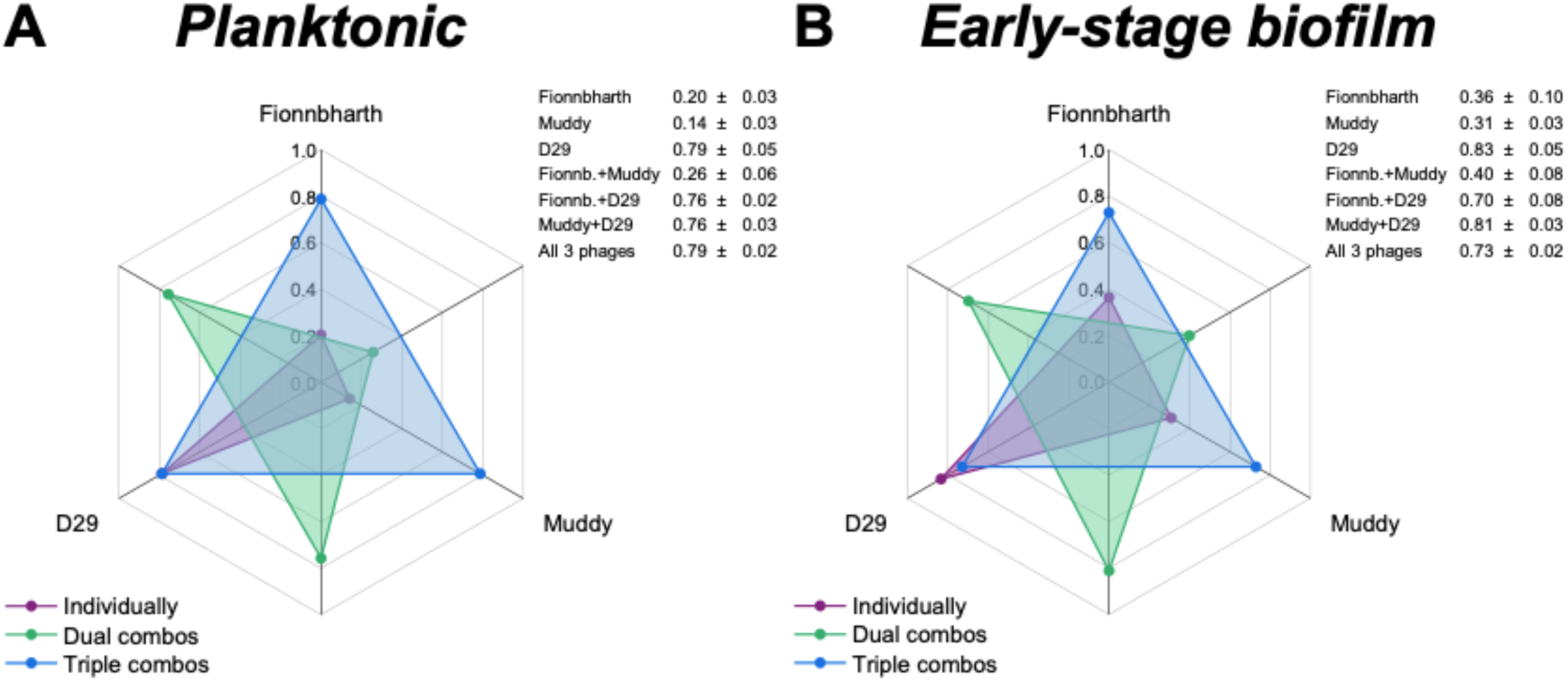
Comparison of the virulence of mycobacteriophages and their cocktail formulations. Spider diagram depicting the virulence index values of phages Fionnbharth, Muddy, D29 and their combinations infecting planktonic (A) and early-stage biofilm (B) *M. smegmatis*. The virulence index scale increases from a minimum of 0 (central point of the chart) to a maximum of 1 (outermost circle). Each axis corresponds to a phage (labelled on the outside of the chart) or their combinations (interposed axes). The virulence index values of the different combinations are indicated by dots in the spider diagram (orange: individual phage; green: two-phage cocktails; red: three-phage cocktail) and listed in the insert tables (average ± standard deviation, n ≥ 3).

**Figure 3.**
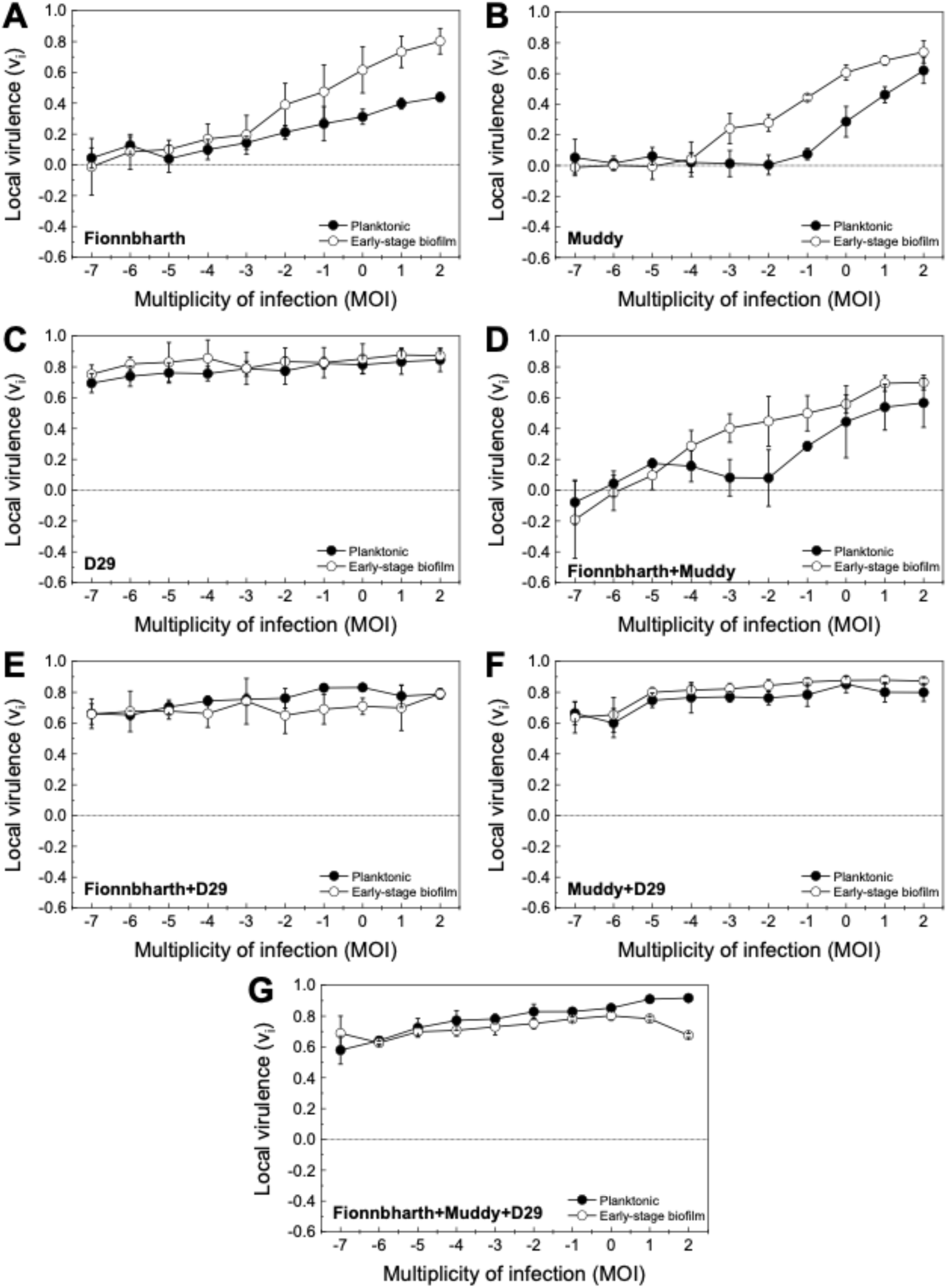
Virulence in planktonic and early-stage biofilm *M. smegmatis*. Virulence curves for phages Fionnbharth (A), Muddy (B), D29 (C), the two-phage cocktails (Fionnbharth + Muddy (D), Fionnbharth + D29 (E), Muddy + D29 (F)), and the three-phage cocktail (Fionnbharth + Muddy + D29 (G)) infecting *M. smegmatis* in planktonic (filled circle) growth conditions. The results are compiled for planktonic (H) and early-stage biofilm (I) for comparison purposes. Values are reported as average ± standard deviation (n ≥ 3).

Similar trends were observed for the phages against the early-stage biofilm phenotype (virulence index: Figure 2B; virulence curves: Figure 3; reduction curves: Supplementary Figure 2). D29 was significantly more virulent than Fionnbharth and Muddy (General Linear Model, P < 0.001; Figure 3I), yielding a higher virulence index value (V_D29-biofilm_ = 0.83, V_Fionn-biofilm_ = 0.36 and V_Muddy-biofilm_ = 0.31; one-way ANOVA, P < 0.001, respectively; Figure 2B). The combination of Fionnbharth and Muddy had a simple additive effect on virulence (Figure 3I). Finally, here again, the infection kinetics of all cocktail combinations containing D29 did not significantly differ from D29 used alone (General Linear Model, P > 0.05; Figure 3I).

### 3.2 Virulence against planktonic and early-stage biofilm phenotypes

Interestingly, the virulence of both Fionnbharth and Muddy increased during infection against early-stage biofilm compared to planktonic phenotype, from MOI 10^-2^ and 10^-3^, respectively (one-way ANOVA, Fionnbharth: P < 0.05, Muddy: P < 0.01; Figure 3A-B), reaching higher virulence index values of V_Fionn-biofilm_ = 0.36 and V_Muddy-biofilm_ = 0.31 (one-way ANOVA, Fionnbharth: P < 0.01, Muddy: P < 0.001; Figure 2). The increase in virulence against early-stage biofilm was similarly observed when Fionnbharth and Muddy were used in combination at MOIs ≥ 10^-4^ (one-way ANOVA, P < 0.05; Figure 3D), with a higher virulence index of V_Fionn+Muddy-biofilm_ = 0.40 as compared to V_Fionn+Muddy-planktonic_ = 0.26 (one-way ANOVA, P < 0.05; Figure 2). One hypothesis is that this may be due to the infection of the early-stage biofilm phenotype leading to more efficient infections, through one or a combination of factors, such as more effective phage adsorption to their host due to the expression of different receptors or different receptor density on the host cell, better binding of the phage receptor binding proteins on the host cell surface, reduced eclipse period, greater burst size, etc. In all cases, this would be consistent with the fact that mycobacteria-phage co-evolution should mostly have taken place in environments similar to the early-stage biofilm growth conditions tested here.

Another factor to consider is that, under this growth condition, *M. smegmatis* grows in agglomerates (microcolonies and biofilm pellicles), where, at high enough phage densities, host-to-host proximity may favor phage predation^43^. Conversely, infection kinetics of D29 did not change between infection of hosts growing as planktonic or early-stage biofilm cultures (one-way ANOVA, P > 0.05; Figure 3C); the same was observed for all two- and three-phage combinations containing D29 (one-way ANOVA, P > 0.05; Figure 3E–G). This suggests that the early-stage biofilm phenotype of *M. smegmatis* can replicate D29 as efficiently and rapidly as the planktonic phenotype.

### 3.3 Adsorption to planktonic M. smegmatis

As adsorption is an important step of phage infection and propagation, it was investigated for the phages used in this study in two ways. The first consisted of assessing adsorption to *M. smegmatis* previously grown as a planktonic culture, with and without shaking (Figure 1). This gives us information on the effects of changes in collision rates and planktonic phenotypes on the adsorption process. The results are shown in Figure 4 (comparison between shaking and static adsorption) and Supplementary Figure 3 (comparison between phages). The resulting adsorption efficiency (ε) and rate constant × initial host concentration (k·C_h0_) values are summarized in Table 1. Under shaking adsorption (Supplementary Figure 3A), D29 showed a rapid decrease in free phage fraction, followed by stabilization to a plateau, indicative of the adsorption efficiency, with no further adsorption after approximately 20 min. This adsorption behavior was consistent with results reported in the literature for this phage^44–46^; Fionnbharth and Muddy showed similar overall trends. However, Fionnbharth exhibited faster adsorption (higher k·C_h0_; two-sample t-test, P < 0.01) and lower final free phages fraction (higher adsorption efficiency; two-sample t-test, P < 0.05) than D29. Muddy, on the other hand, displayed slower (lower k·C_h0_; two-sample t-test, P < 0.01 for both D29 and Fionnbharth) and less efficient (lower adsorption efficiency; two-sample t-test, P < 0.01 and P < 0.001 for D29 and Fionnbharth, respectively) adsorption than the other two phages.

**Figure 4.**
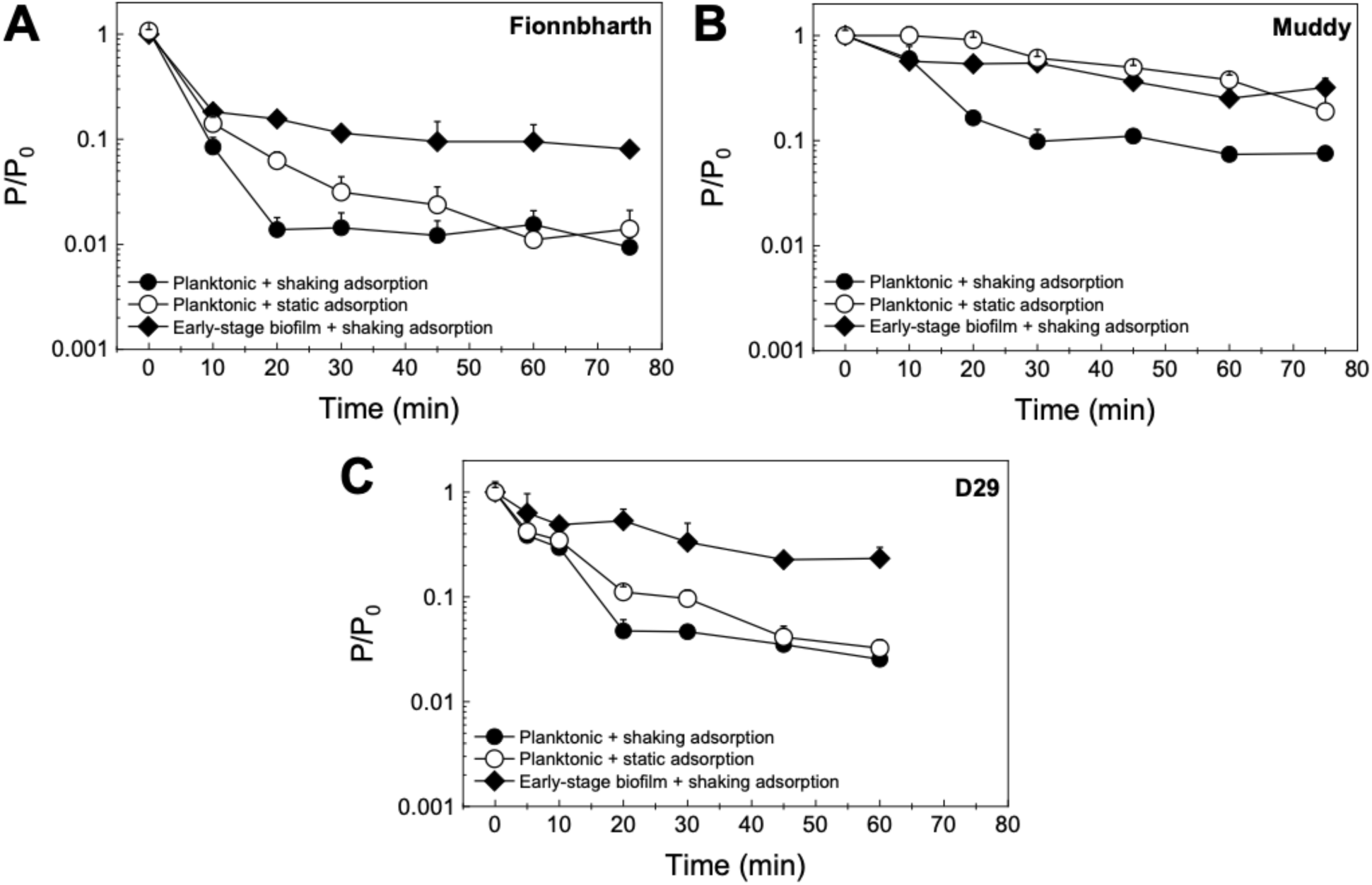
Adsorption to *M. smegmatis*. The free phage concentration P relative to the initial phage concentration P_0_ of Fionnbharth (A), Muddy (B), and D29 (C) adsorbing to planktonic or early-stage biofilm *M. smegmatis*. Adsorption to the planktonic phenotype was measured under shaking (filled circle) and static (empty circle) conditions; adsorption to early-stage biofilm phenotype was measured under shaking conditions (filled diamonds). All data is shown as average ± standard deviation, n ≥ 3).

**Table 1.**
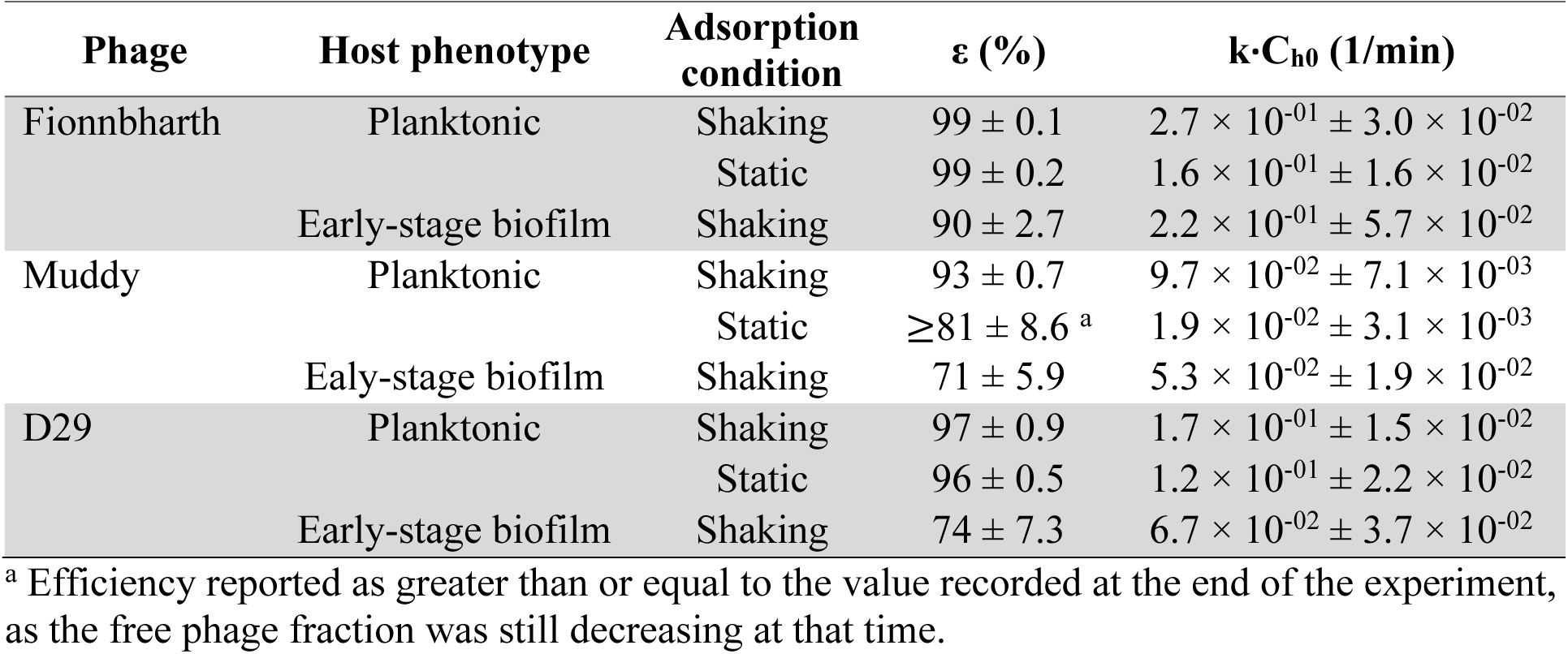
Adsorption parameters (efficiency (ε) and rate constant × initial host concentration (k·C_h0_)) calculated for phages Fionnbharth, Muddy and D29 adsorbing to different *M. smegmatis* phenotypes (planktonic and early-stage biofilm) under shaking and static conditions (average ± standard deviation, n = 3–4).

Similar comparisons between the three phages can be made under static adsorption (Figure 4, Supplementary Figure 3B), where Fionnbharth had the fastest adsorption, followed by D29 and then Muddy (two-sample t-test, Fionnbharth vs Muddy P < 0.001, Fionnbharth vs D29 P < 0.05, D29 vs Muddy P < 0.01). Although little is known about the receptors of these three mycobacteriophages^47^, it appears many preferentially bind to sections of cell wall synthesis at the poles and septa of mycobacteria^48^. Here, we can hypothesize that the differences in adsorption kinetics, efficiencies, and trends may be due to the phages targeting different receptors. It is also worth noting that, although D29 displayed the highest virulence, it did not exhibit the fastest adsorption or highest adsorption efficiency, highlighting the importance of other factors (e.g., eclipse period, latent period, burst size) on the overall infection process.

Looking at the differences in adsorption behavior under shaking and static conditions, all three phages were slower to adsorb to planktonic *M. smegmatis* under static than under shaking conditions (two-sample t-test, Fionnbharth P < 0.01, Muddy P < 0.001, D29 P < 0.05) (Figure 4). The results observed are consistent with the fact that, under shaking, there is an increase in collision rate, leading to more opportunities for initial contact between phage and host per unit time; therefore, the phages displayed faster adsorption kinetics.

### 3.4 Adsorption to early-stage biofilm of M. smegmatis

The adsorption tests were also conducted following a second approach: *M. smegmatis* was grown as early-stage biofilms before assessing adsorption under shaking conditions (Figure 1). By comparing these results with those obtained on planktonic *M. smegmatis*, it is possible to see the effect of the phenotype of the host on the adsorption behavior of the phages. The results are shown in Figure 4 (comparison between planktonic and early-stage biofilm *M. smegmatis*) and Supplementary Figure 4 (comparison between phages). The ε and k·C_h0_ values are summarized in Table 1. For D29, the early-stage biofilm phenotype leads to significantly slower adsorption than the planktonic phenotype (two-sample t-test, P < 0.05) (Figure 4C). Additionally, all three phages exhibited lower adsorption efficiencies to the early-stage biofilm phenotype than to the planktonic phenotype under shaking conditions (two-sample t-test, Fionnbharth P < 0.01, Muddy P < 0.01, D29 P < 0.05) (Figure 4).

The slower adsorption kinetics (shown through lower k·C_h0_ values) and lower adsorption efficiencies suggest a reduced availability of receptors in the early-stage biofilm of *M. smegmatis*, caused either by the increased expression of the extracellular matrix of mycobacterium (leading to larger aggregates and pellicle formation) or by a reduced surface density of the receptors on the cell surface in this phenotype.

However, it is also important to note that, despite the reduced adsorption of the phages infecting the early-stage biofilm host phenotype (Table 1), the virulence of Fionnbharth and Muddy–and their 2-phage cocktail–actually increased in infections performed against hosts grown as early-stage biofilm, and the virulence of D29–and the virulence of cocktails containing D29–did not decrease (Figure 2 and Figure 3) . This suggests that, despite the observed reduced adsorption, other parameters in the infection process (e.g., shorter eclipse time, shorter lysis time, larger bust size, proximity of host cells in agglomerates, etc.) are favored in the early-stage biofilm.

Finally, when comparing the adsorption of the three mycobacteriophages to the early-stage biofilm (Figure 4), Muddy and D29 displayed similar adsorption rate and efficiency (two-sample t-test, P > 0.05 for both). Fionnbharth had the fastest adsorption rate (two-sample t-test, P < 0.05 for both Muddy and D29), which is a similar trend to what was observed when adsorbing to the planktonic phenotype (Figure 4, Supplementary Figure 3A).

These comparisons suggest that the three mycobacteriophages may target different receptors; however, future studies on the receptors of these phages will be needed to confirm this.

## 4. Conclusions

Characterization of infection dynamics of mycobacteriophages is an important step towards development and optimization of therapies and other technologies to counteract TB and nontuberculous mycobacteria infections. The virulence of three clinically relevant mycobacteriophages was studied by applying the virulence index method to the infection of planktonic and early-stage biofilm cultures of *M. smegmatis*. Phage D29, used both individually and in two-phage and three-phage cocktails, had the highest virulence. One interesting observation was that phages Fionnbharth and Muddy, as well as their two-phage cocktail, had higher virulence against the early-stage biofilm host phenotype than the planktonic culture phenotype. This points towards these two phages potentially being better adapted to infecting early and established biofilms—which are more representative of their natural environment. Another possibility is that host-to-host proximity in biofilms favors the rapid propagation of the infection. This could be advantageous for therapies, as these phages seem more efficient in environments similar to the conditions encountered during mycobacterial infection. Finally, an important step in the infection process, adsorption, was also compared under similar conditions. Interestingly, despite displaying the highest virulence among the three phages, D29 did not exhibit the fastest or most efficient adsorption. Additionally, adsorption kinetics and efficiency were generally higher against the planktonic cultures than against the early-stage biofilm, despite the three phages and their cocktail formulations being either as or more virulent against the early-stage biofilm phenotype. These observations underscore the contribution of other infection parameters in establishing virulence. Overall, the results of this comparative study inform us on mycobacteriophage infections under different host growth conditions (planktonic and early-stage biofilm cultures), and provide crucial information on infection dynamics for cocktail formulations and eventual treatment strategies.

## Authorship contributions

**Ilaria Rubino:** Conceptualization, Methodology, Validation, Formal analysis, Investigation, Data Curation, Writing - Original Draft, Visualization. **Carlos A. Guerrero:** Methodology, Resources, Writing - Review & Editing. **Melissa Harrison:** Validation, Investigation, Data Curation, Writing - Review & Editing. **Sheila Co:** Methodology, Writing - Review & Editing. **Isobel Tetreau:** Methodology, Writing - Review & Editing. **Mani Ordoubadi:** Methodology, Writing - Review & Editing. **Sasha E. Larsen:** Methodology, Writing - Review & Editing, Funding acquisition. **Rhea N. Coler:** Methodology, Writing - Review & Editing, Funding acquisition. **Reinhard Vehring:** Methodology, Writing - Review & Editing, Supervision, Funding acquisition. **Graham F. Hatfull:** Methodology, Resources, Writing - Review & Editing, Supervision, Funding acquisition. **Dominic Sauvageau:** Conceptualization, Methodology, Resources, Writing - Original Draft, Visualization, Supervision, Project administration, Funding acquisition.

## Acknowledgments

This research was financially supported by the Alberta Innovates Accelerating Innovations into CarE program(Sauvageau), the Natural Sciences and Engineering Research Council of Canada Discovery grant program (Sauvageau), the Claire-Deschênes Postdoctoral Fellowship (Rubino), the National Institute of Allergy and Infectious Diseases of the National Institutes of Health grant AI156807-01 (Coler and Larsen), and the National Institutes of Health grant GM131729 (Hatfull) and Howard Hughes Medical Institute grant GT2053 (Hatfull).

## Supplementary Data

Provided with this manuscript.

## Data availability

All data generated or analyzed during this study are included in this published article (and its supplementary information files).

