## Supplemental Figures for "Comparative study on the virulence of mycobacteriophages"

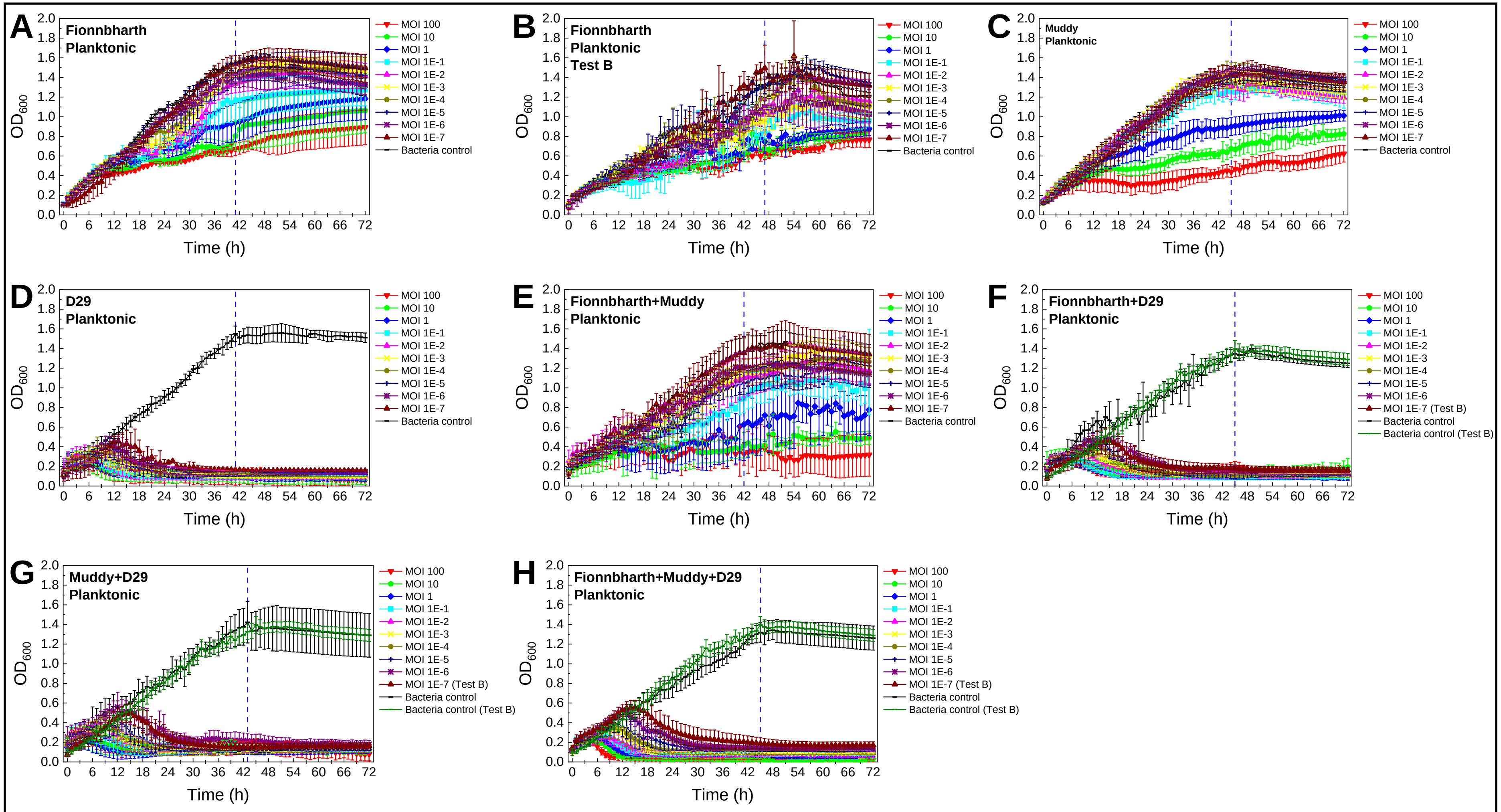


**Supplementary Figure 1. Reduction curves for infection under planktonic growth.** Reduction curves for phages Fionnbharth (A,B), Muddy (C), D29 (D) and the two-phage (E–G) and three-phage (H) cocktails infecting *M. smegmatis* at MOIs $10^{-7}$–100 under planktonic growth (average ± standard deviation, n ≥ 3).

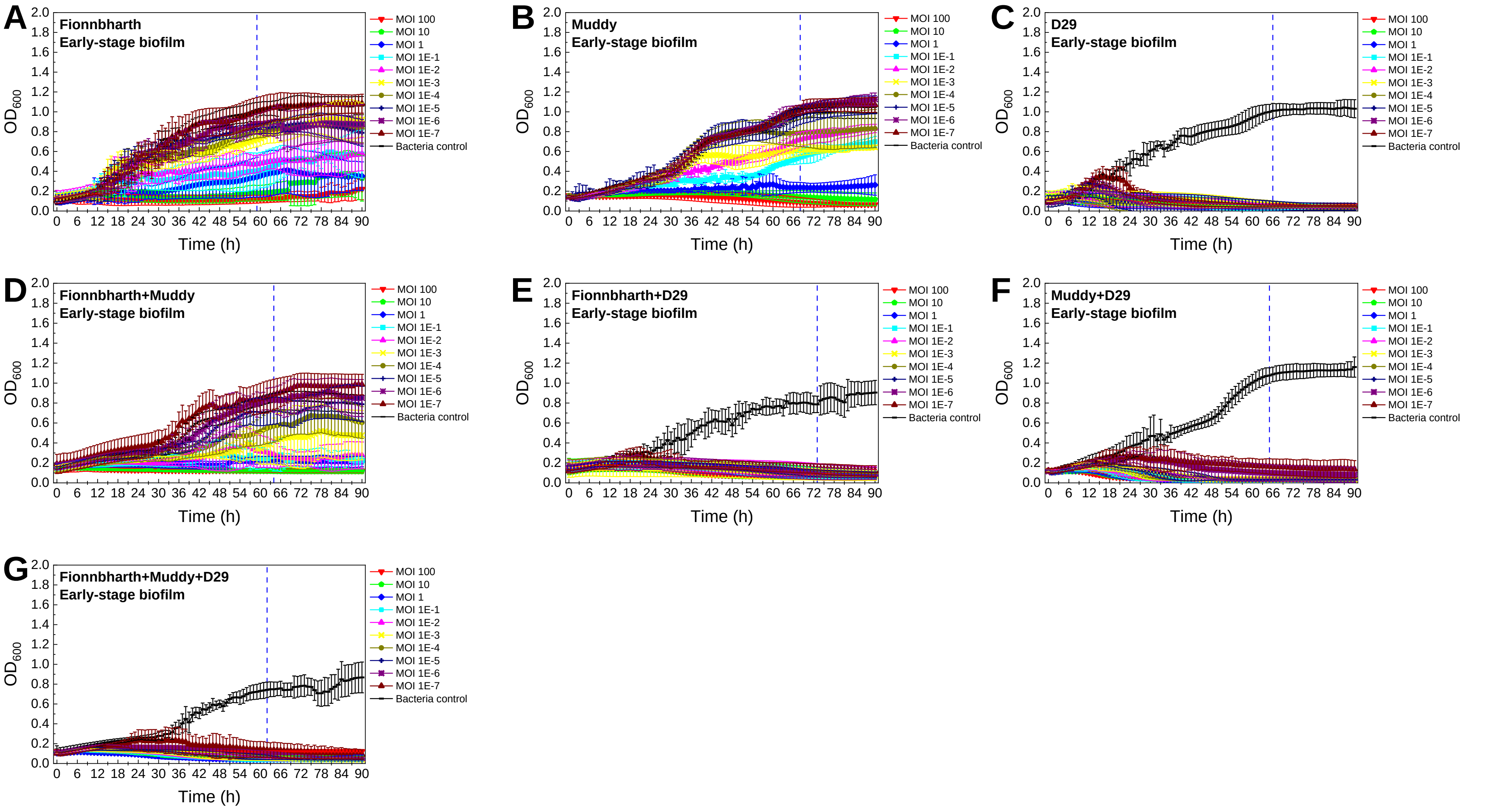


**Supplementary Figure 2. Reduction curves under early-stage biofilm growth.** Reduction curves for phages Fionnbharth (A), Muddy (B), D29 (C) and the two-phage (D–F) and three-phage (G) cocktails infecting *M. smegmatis* at MOIs $10^{-7}$–100 under early-stage biofilm growth (average ± standard deviation, n ≥ 3).

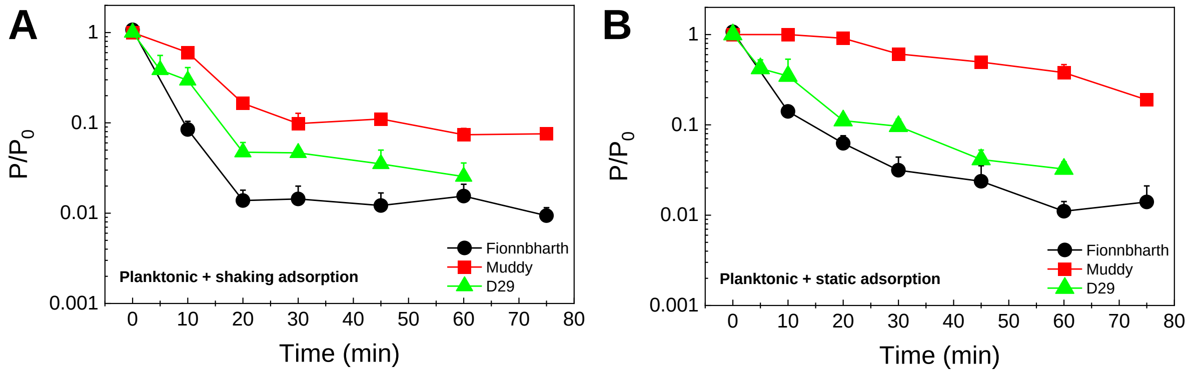


**Supplementary Figure 3. Comparison of the adsorption of mycobacteriophages to planktonic *M. smegmatis*.** The free phage concentration P relative to the initial phage concentration $P_0$ of Fionnbharth, Muddy, and D29 adsorbing to planktonic *M. smegmatis*. The adsorption was measured under shaking (A) and static (B) adsorption conditions (average ± standard deviation, n ≥ 3).

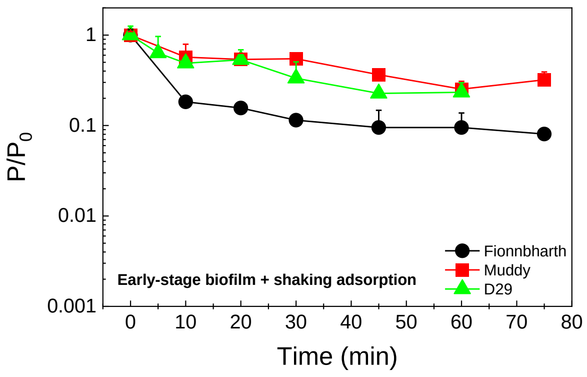


**Supplementary Figure 4. Comparison of the adsorption of mycobacteriophages to early-stage biofilm of *M. smegmatis*.** The free phage concentration P relative to the initial phage concentration $P_0$ of Fionnbharth, Muddy, and D29 adsorbing to early-stage biofilm of *M. smegmatis*. The adsorption was measured under shaking adsorption conditions (average ± standard deviation, n ≥ 3).
